# Selection of Pairings Reaching Evenly Across the Data (SPREAD): A simple algorithm to design maximally informative fully crossed mating experiments

**DOI:** 10.1101/009720

**Authors:** Kolea Zimmerman, Daniel Levitis, Ethan Addicott, Anne Pringle

## Abstract

We present a novel algorithm for the design of crossing experiments. The algorithm identifies a set of individuals (a “crossing-set”) from a larger pool of potential crossing-sets by maximizing the diversity of traits of interest, for example, maximizing the range of genetic and geographic distances between individuals included in the crossing-set. To calculate diversity, we use the mean nearest neighbor distance of crosses plotted in trait space. We implement our algorithm on a real dataset of *Neurospora crassa* strains, using the genetic and geographic distances between potential crosses as a two-dimensional trait space. In simulated mating experiments, crossing-sets selected by our algorithm provide better estimates of underlying parameter values than randomly chosen crossing-sets.

## INTRODUCTION

Researchers planning mating experiments are faced with a critical design choice— deciding how many pairs and which pairs of individuals to mate. The number of crosses in a mating experiment can influence statistical estimates of genetic effects and combining abilities (Jui and Lefkovitch, 1992). The selection of pairs to use in a mating experiment also affects the outcome of the experiment. For example, if the goal of a mating experiment is to understand the genetic basis of a trait, as in quantitative trait locus (QTL) analysis, then parents should carefully be chosen to maximize genetic diversity among offspring and increase the likelihood of detecting QTLs (Crepieux *et al.*, 2004). The increasing accessibility of population genetic and genomic datasets offer genetic data on more individuals than can reasonably be used in most experiments (Cushman, 2014). This poses a methodological problem: how to choose a subsample of mating pairs that best reflects the range of cross characteristics (in two or more dimensions of genetic, geographic, or ecological space) of the complete set of all available pairs.

One solution is to select a subsample that recapitulates characteristics of the larger set and preserves underlying relationships between the variables used to define a trait space. The representative subsample might mimic the broad distribution of crosses in the larger set, in other words, it attempts to maintain the shape, clumps, etc. of the larger set. Other subsampling methods include choosing samples by eye or randomly, but these methods may truncate the trait space by omitting outliers or disproportionately drawing from the dense center of a distribution. Omissions in sampling may hinder a complete understanding of how response variables, for example reproduction, vary across the trait space of all possible crosses. Furthermore, predicting response variables outside the range of explanatory variables used in an experiment involves extreme value methods, which can increase the error associated with predictions, unless limiting assumptions are made (Pauli and Coles, 2001).

In breeding and mating studies, fully crossed matings between all possible pairs are desirable to determine combining ability and genetic and maternal effects (Griffing, 1956); (Zhu and Weir, 1996). But a directed subsampling of all potential crosses to achieve some other aim, for example to maximize genetic diversity within the experiment, may result in a set of target individuals that cannot be fully crossed. A method is required to subsample and design matings to satisfy both aims, for example to maximize genetic diversity and to fully cross included individuals.

Algorithms for maximizing combinatorial diversity have been extensively developed in the context of generating diverse molecular libraries for drug screening (Martin and Critchlow, 1999). In these algorithms the metric of diversity is based on “redundancy” and “coverage” (Martin and Critchlow, 1999). Redundancy is the overlapping or clumping of points in space, while coverage is the spread of points across the space. An ideal diversity metric would minimize redundancy while maximizing coverage. The algorithms used in chemical combinatorial analysis focus on maximizing the diversity of a subset of molecules from a larger set by step-wise analysis of differences between additional compounds added to a set (Holliday *et al.*, 1995). These algorithms cannot be directly applied to our problem because they do not require selection of fully crossed sets. However, we use their definitions of ideal set diversity to derive our own measure of diversity that can be applied to fully crossed sets.

Calculating the mean of the nearest neighbor distances (NND) of points representing a full factorial set of crosses plotted based on their underlying parameters (*e.g*., genetic, geographic or ecological distance) will give a measure of the evenness or “non-redundancy” of the points. The mean NND is often used to determine if a particular set of plotted points is randomly distributed or not (Clark and Evans, 1954). A set of plotted points that are clumped will result in a smaller value of the mean NND than a sample with the same number of more evenly and broadly distributed points. The maximum mean NND (MMNND) will occur when points are spread as evenly as possible and the “coverage” of space is maximal (Wang and Cumming, 2011). Thus, identifying a set of crosses with the MMNND from a large random sample of many potential sets of crosses (“crossing-sets”) will return a crossing-set that is both broad and even with respect to underlying trait values as compared to a randomly sampled crossing-set.

We introduce a simple algorithmic sampling method for choosing crossing-sets; we name the algorithm SPREAD (Selection of Pairings Reaching Evenly Across the Data). SPREAD is based on selecting the single crossing-set with the MMNND from among a large random sample of potential crossing-sets plotted on two-dimensional trait space. We use our algorithm to select a crossing-set from a genotyped collection of geographically widespread wild strains of the filamentous fungus, *Neurospora crassa.* Strains of this fungus have one of two mating types, denoted mat-A or mat-a. The two parents in a cross must have different mating types to mate. Recently, 24 strains of each mating type were genotyped using RNAseq (Ellison *et al.*, 2011). The genotyped strains were collected from diverse locations, allowing us to assign both genetic (the number of different SNPs) and geographic (the distance between collection sites) distance values to each of the 576 potential crosses. Using this dataset as our example, we implemented the SPREAD algorithm and tested the effectiveness of the SPREAD algorithm when the true MMNND is not easily calculable. Finally, we compare the ability of SPREAD selected and randomly sampled crossing-sets to estimate known parameter values that relate genetic and geographic distances to reproductive output for the entire set of all potential crosses.

## METHODS

### Description of the SPREAD Algorithm

Define 𝕏 and 𝕐 as the set of available strains or individuals of each mating type or sex (‘type’) *x* and *y* respectively, and *s*_*x*_ × *s*_*y*_ as the feasible number of crosses that can be completed in an experiment. The variables *s*_*x*_ and *s*_*y*_ are the number of strains selected for the experiment and are less than |𝕏| and |𝕐| respectively. Draw a large number, *h*, of random samples containing *s*_*x*_ and *s*_*y*_ strains of each type from all possible sets of strains 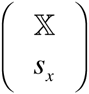 and 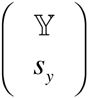. For each of the *h* samples, plot crosses based on values associated with the crosses (for example number of differing SNPs vs. geographic distance), and then calculate the mean of the NNDs of all plotted crosses. Generate a list of *h* mean NNDs. Finally, use the maximum value from the list because it corresponds to the set of *s*_*x*_ × *s*_*y*_ strains that most broadly and evenly represents the parameter of interest. A formal mathematical description of this algorithm is available at http://dx.doi.org/10.6084/m9.figshare.1180170.

### A worked example using SPREAD

We used a previously published population genomics dataset consisting of single nucleotide polymorphisms (SNPs) from transcriptomes of geographically diverse wild isolates of the fungus *N. crassa* to test our method (Ellison *et al.*, 2011). We started with the set of all pairwise combinations of strains and then filtered to include only mating type compatible pairs. We calculated genetic distances between compatible pairs by counting the number of different SNPs between each pair and calculated geographic distances using the great-circle distance between strain locales. The genetic and geographic distance values for each pair were used to map all the crosses on genetic and geographic distance axes. This is the “original distribution” of crosses.

We randomly sampled *h* = 1000 lists of *s*_*A*_ = *s*_*a*_ = 12 strains of each mating type from the set of all 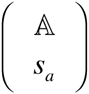 and 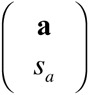 strains without replacement, where 𝔸 and **a** and are the sets of strains available for each mating type; in this case |𝔸| = |**a**| = 24. We computed all possible pairwise mating combinations for each of the 1000 random samples of *s*_*A*_ = *s*_*a*_ = 12 strain lists, resulting in 1000 crossing-sets each containing 144 crosses. We then plotted each crossing-set on geographic vs. genetic distance space and computed the mean nearest neighbor distances using Euclidean distance calculations. The crossing-set with the MMNND of all 1000 crossing-sets was selected. In this worked example, selecting from a random sample of 1000 crossing sets— and not selecting from all possible crossing sets—is necessary because the total number of possible crossing sets in this case is 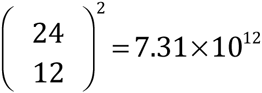. Computing nearest neighbour distances for all possible crossing-sets is computationally prohibitive and, as shown below, unnecessary.

We implemented our algorithm and additional analyses in the R programming language (R Core Team, 2014). R code for the implementation of the SPREAD algorithm on crossing-sets with two traits and two sexes is available at http://dx.doi.org/10.6084/m9.figshare.1180165. The following R packages were used in this analysis: plyr (Wickham, 2011), reshape2 (Wickham, 2007), ggplot2 (Wickham, 2009), spatstat (Baddeley and Turner, 2005), foreach (Weston and Analytics, 2014), and glmmADMB (Fournier *et al.*, 2012; Skaug *et al.*, 2013).

### Evaluating SPREAD’s approximation of the true MMNND

The true MMNND can only be determined if all possible crossing-sets for a given *s*_*A*_ and *s*_*a*_ are evaluated. Therefore, calculating the true MMNND may not be possible, even with high performance computing resources. For example, if M = 300 and F = 300 and a crossing-set is desired with 20 individuals of each type, then the total number of possible crossing-sets would be 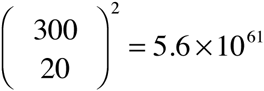. Using a random sample of all available crossing-sets to estimate the MMNND is more practical, especially if the estimated MMNND approximates the true MMNND.

We implemented the algorithm as described above for the *N. crassa* dataset, except we varied crossing-set size by implementing SPREAD for *s*_*A*_ = *s*_*a*_ = 2, 3, 4, …, 22. To simplify the process, we used crossing-sets where *s*_*A*_ = *s*_*a*_, but this is not a requirement of the SPREAD algorithm. We used five different *h* values (1, 10, 100, 1000, and 10000) to compare the effects of *h* size on the MMNND values returned from SPREAD. We included an *h* value of 1 to simulate the distribution of MNND values of SRS-generated crossing-sets. We repeated this process 1000 times to obtain bootstrapped distributions of MMNND values for the different crossing-set sizes and *h* values.

### Comparing model fits of SPREAD- and SRS-generated crossing-sets

Using SPREAD to design fully crossed mating experiments may be more effective than selecting crossing-sets at random because broad and even sampling will provide greater statistical power to understand how dependent variables vary based on cross characteristics (*e.g*., how reproductive success depends on the genetic or geographic distances between parents). To evaluate this hypothesis empirically, we created a simulated dataset of cross outcomes (*i.e*. reproduction) and modeled relationships between reproduction and the characteristics of crosses in crossing-sets generated from SPREAD *versus* those generated by simple random sampling. Simulated experimental data take the form of total ascospore counts (the sexually produced spores of the fungus).

First, we generated simulated data for all possible crosses of the entire crossing-set of 24 mat-A × 24 mat-a strains, using a generalized linear model (GLM) fitted to unpublished empirical data. This model was evaluated using the glmmADMB package (Fournier *et al.*, 2012; Skaug *et al.*, 2013) as *Total Ascospore Count* = *Genetic Distance* + *Geographic Distance* + (*Genetic Distance*)^2^ + (*Geographic Distance*)^2^ + *Genetic Distance : Geographic Distance*. The response variable of *Total Ascospore Count* was modeled with a negative binomial distribution using a log link function. Genetic and geographic distance values were mean-centered to prevent autocorrelation between linear and quadratic parameters. We did not include all possible interaction terms because we wanted to restrict the model to only linear and quadratic terms. Our choice is based on the biological hypothesis that there is a single optimum of reproductive output, intermediate between inbreeding and outbreeding depression (Lynch, 1991).

Second, we simulated four experimental replicates for each possible cross by drawing from a negative binomial distribution with a mean derived from the predicted experimental values and the negative binomial dispersion parameter derived from the empirical data model. “True” model parameter values were determined by evaluating the model described above with the complete simulated data set of all crosses.

Using the complete set of simulated experimental data we computed data sets for 1000 different crossing-sets generated with either SPREAD or SRS. The algorithm parameter values for the SPREAD-generated crossing-sets were *s*_*A*_ = *s*_*a*_ = 12 and *h* = 1000. We chose *s*_*A*_ = *s*_*a*_ = 12 to test the edge case of a maximally complex sample space (the largest number of possible crossing-set permutations occurs when *s*_*A*_ = *s*_*a*_ = 12). Crossing-sets chosen by SRS were of the same size. Model fits were computed for each crossing-set using the model described above. Parameter values and standard errors of the parameter values were recorded for each of the 1000 SPREAD- or SRS-generated crossing-sets. We recorded the MNND values of all SPREAD- and SRS-generated crossing-sets used in this comparison to determine the relationship between MNND of crossing-sets and the ability of those crossing-sets to estimate the true parameter values of the entire population.

## RESULTS

### The worked example

We used SPREAD on the *N. crassa* dataset described above to select a crossing-set with 12 mat-A and 12 mat-a strains. A graphical assessment of the chosen crossing-set plotted on geographic and genetic distance axes shows that our method produces a crossing-set that broadly and evenly represents all potential crosses (Figure 1).

**Figure 1.**
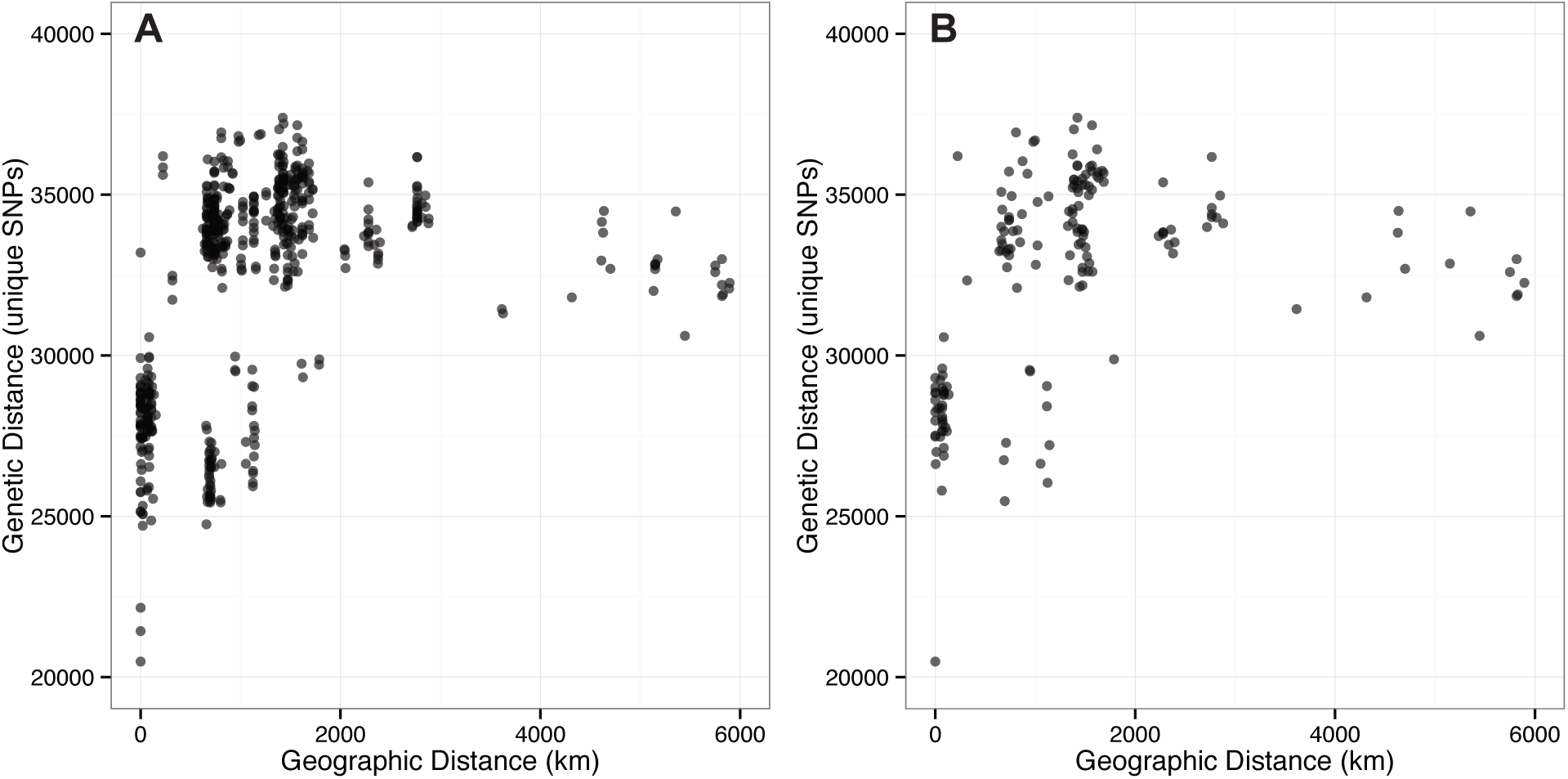
Example implementation of SPREAD. Panel **A** shows all possible crosses between 24 mat-A and 24 mat-a *N. crassa* strains plotted based on genetic and geographic distance between the parents in a cross; panel **B** shows the set of 12 mat-A × 12 mat-a *N. crassa* strains returned from SPREAD (with *h* = 1000). Crosses are plotted as semitransparent dots and darker colors mark overlapping crosses.

### Implementing SPREAD without knowing the true MMNND

While the goal of SPREAD is to find a crossing set with a high MNND value, finding the true maximum MNND is not necessary for most experimental purposes. Rather, the number of randomly selected potential crossing sets considered (*h*) should balance the desirability of finding a set with a high MNND with computational convenience. Plotting distributions of MMNND values for the five *h* values shows that *h* values beyond 1000 deliver sharply diminishing marginal returns for all crossing-sets with more than 16 crosses (Figure 2). Furthermore, the range of the MMNND estimates decreases as *h* increases, mainly driven by an increase of the lower bound while the upper bound remains relatively unchanged.

**Figure 2.**
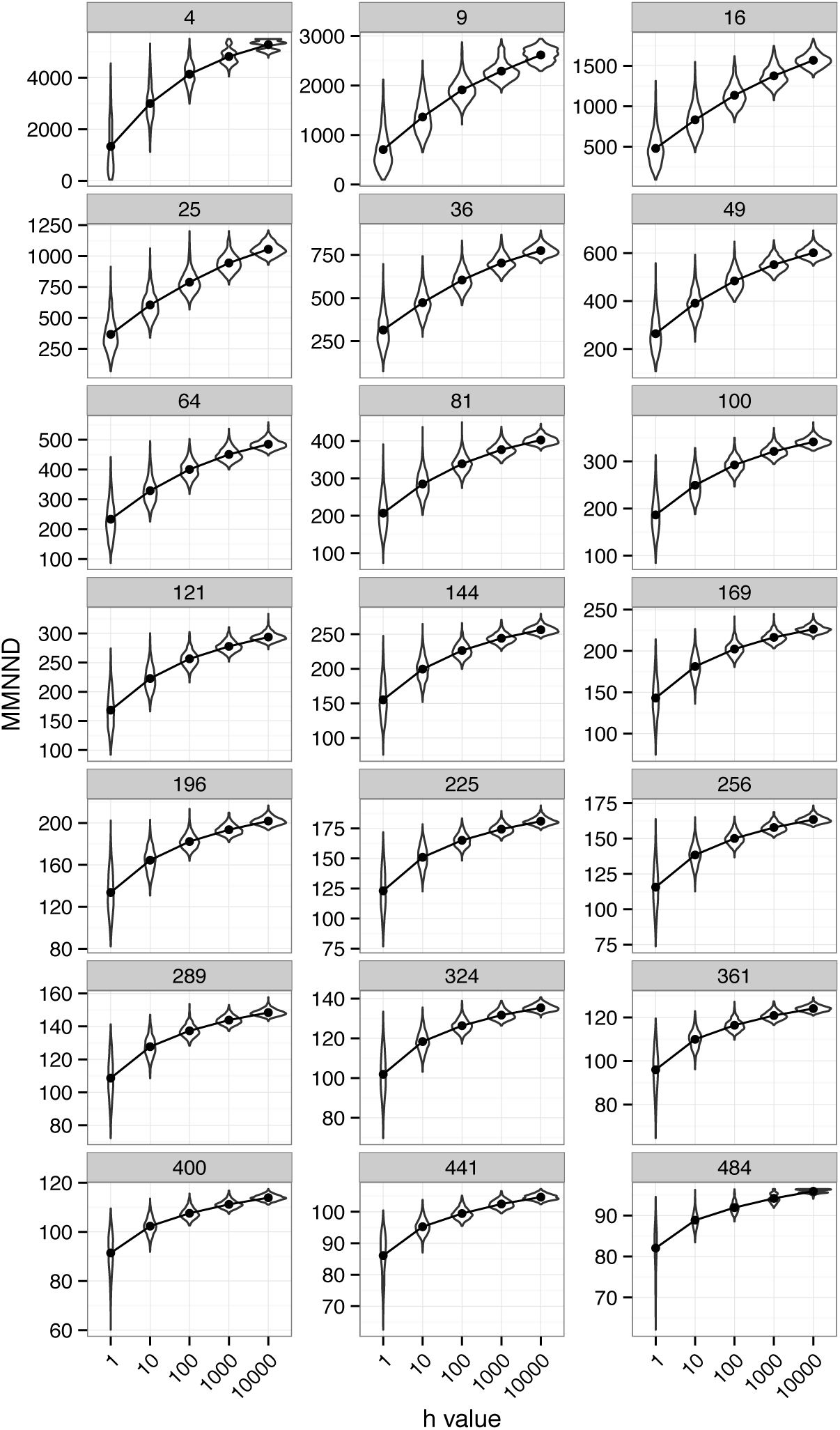
Effect of crossing-set size and *h* value (the number of random crossing-sets from which the crossing-set is selected) on distributions of MMNND values of selected crossing-sets. Each facet is labeled with the size of the crossing-set. With larger *h*-values, estimates of MMNND values increase. As crossing-set size increases MMNND values decrease because with more crosses the average distance between crosses decreases. Violin plots show the entire distributions of MMNND values. Points and lines show how the mean of the distributions changes with increasing *h* values.

### Comparing SPREAD to SRS

We tested the ability of SPREAD to generate crossing sets that will more accurately predict the parameter values one might find if all potential matings of an available breeding population were used in an experiment. Repeated simulations yielded distributions of parameter estimates for each model parameter using each method (SPREAD or SRS) (Figure 3). For both SPREAD and SRS, the peaks of these distributions of estimates do not perfectly align with the underlying values. However, the estimates from models using SPREAD are more accurate than those using SRS (Figure 3A and Table 1). The standard errors of estimates from SPREAD-based models are small compared to those from SRS based models (Figure 3B and Table 2), again increasing the probability of closely approximating true parameter values.

**Figure 3.**
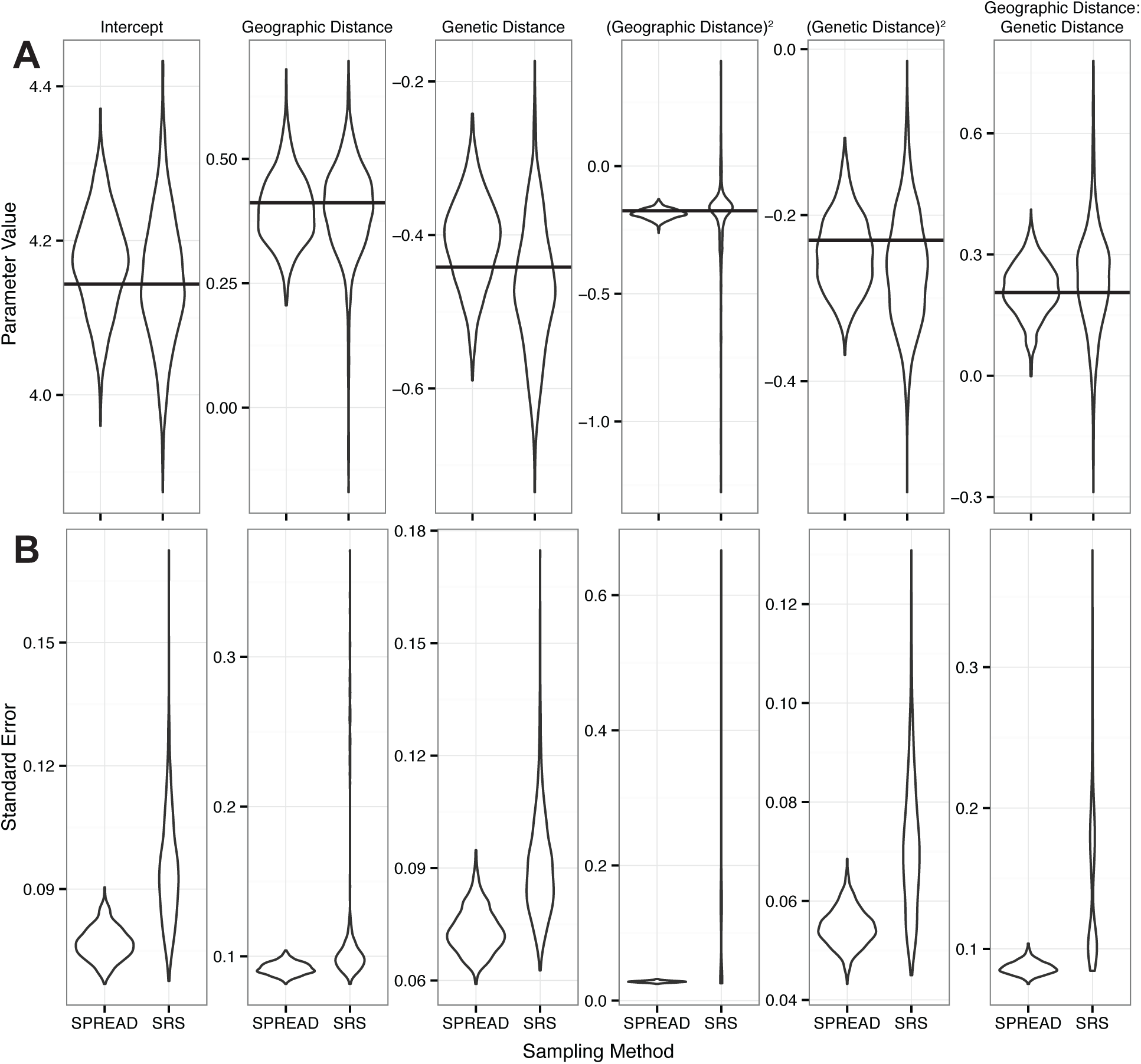
Comparisons of estimated parameter values and standard errors for 1000 crossing-sets generated either with SPREAD or SRS. The SPREAD parameters used are *s*_*A*_ = *s*_*a*_ = 12 and *h* = 1000. Panel **A**: Violin plots showing the distribution of parameter values for each parameter in the model. Horizontal lines indicate the “true” parameter values of the entire simulated *s*_*A*_ = *s*_*a*_ = 24 dataset. Panel **B**: Violin plots showing the distributions of standard errors of parameter values for each parameter in the model.

**Table 1.**
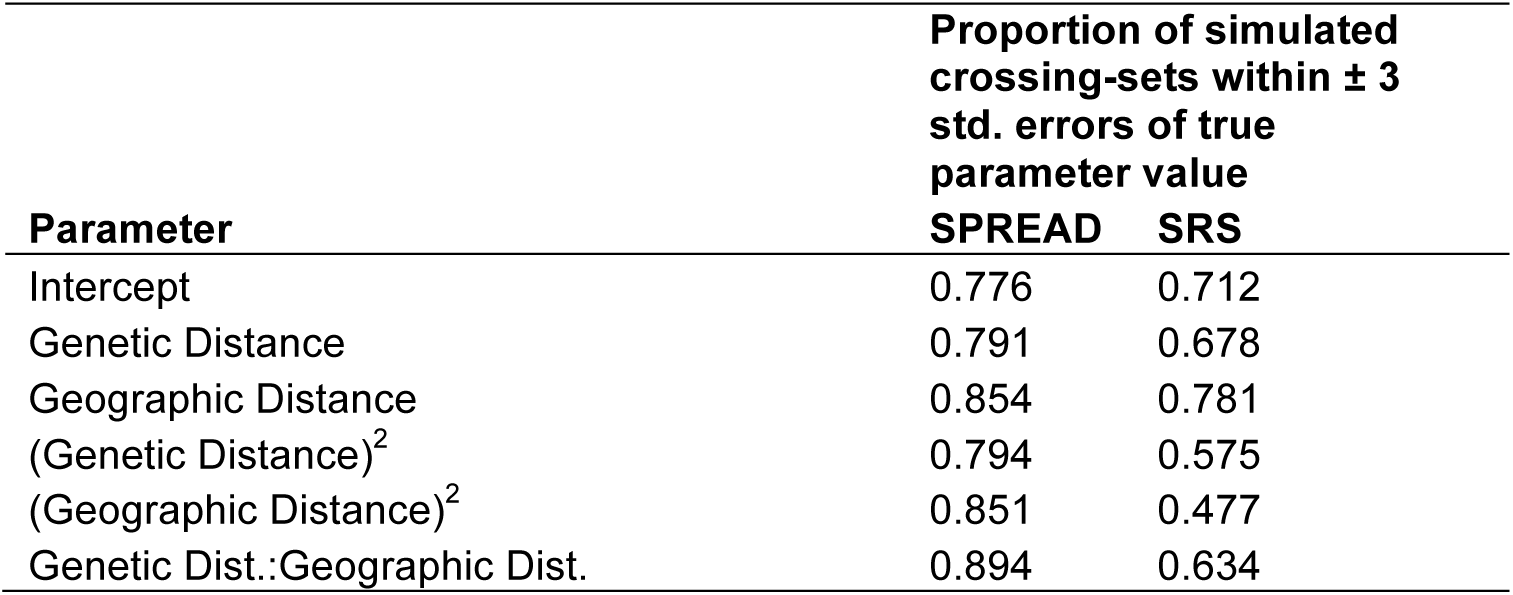
Proportion of the 1000 simulated crossing-sets with parameter values within the range defined by the true parameter value ± 3 standard errors of true parameter values. “Dist.” is the abbreviation for “Distance”.

**Table 2.** Summary statistics for the distribution of standard errors from 1000 model fits using either SPREAD- or SRS-generated crossing-sets. “Dist.” is the abbreviation for “Distance”.

| Parameter | Mean of Std. Errors |  | Variance of Std. Errors |  |
| --- | --- | --- | --- | --- |
|  | SPREAD | SRS | SPREAD | SRS |
| Intercept | 0.0768 | 0.0944 | 1.60E-05 | 1.72E-04 |
| Genetic Distance | 0.0735 | 0.0904 | 3.59E-05 | 1.94E-04 |
| Geographic Distance | 0.0919 | 0.1093 | 1.66E-05 | 1.60E-03 |
| (Genetic Distance)<sup>2</sup> | 0.0547 | 0.0709 | 1.48E-05 | 1.85E-04 |
| (Geographic Distance)<sup>2</sup> | 0.0280 | 0.1079 | 1.79E-06 | 1.19E-02 |
| Genetic Dist.:Geographic Dist. | 0.0867 | 0.1466 | 2.16E-05 | 2.87E-03 |

### MNND vs. parameter estimates

Broadly and evenly distributed explanatory variables increase the accuracy and precision of predictions based on those variables. The MNND is a measure of the broadness and evenness of points in space, and so it should be negatively correlated with measures of inaccuracy or error. Indeed, we found strong negative relationships between the MNND of crossing-sets and both the deviation from true parameter values (Figure 4A) and the standard errors of parameters (Figure 4B). Within SRS-generated crossing-sets, those sets with higher MNND values also recapitulated the true parameter values better than crossing-sets with lower MNND values.

**Figure 4.**
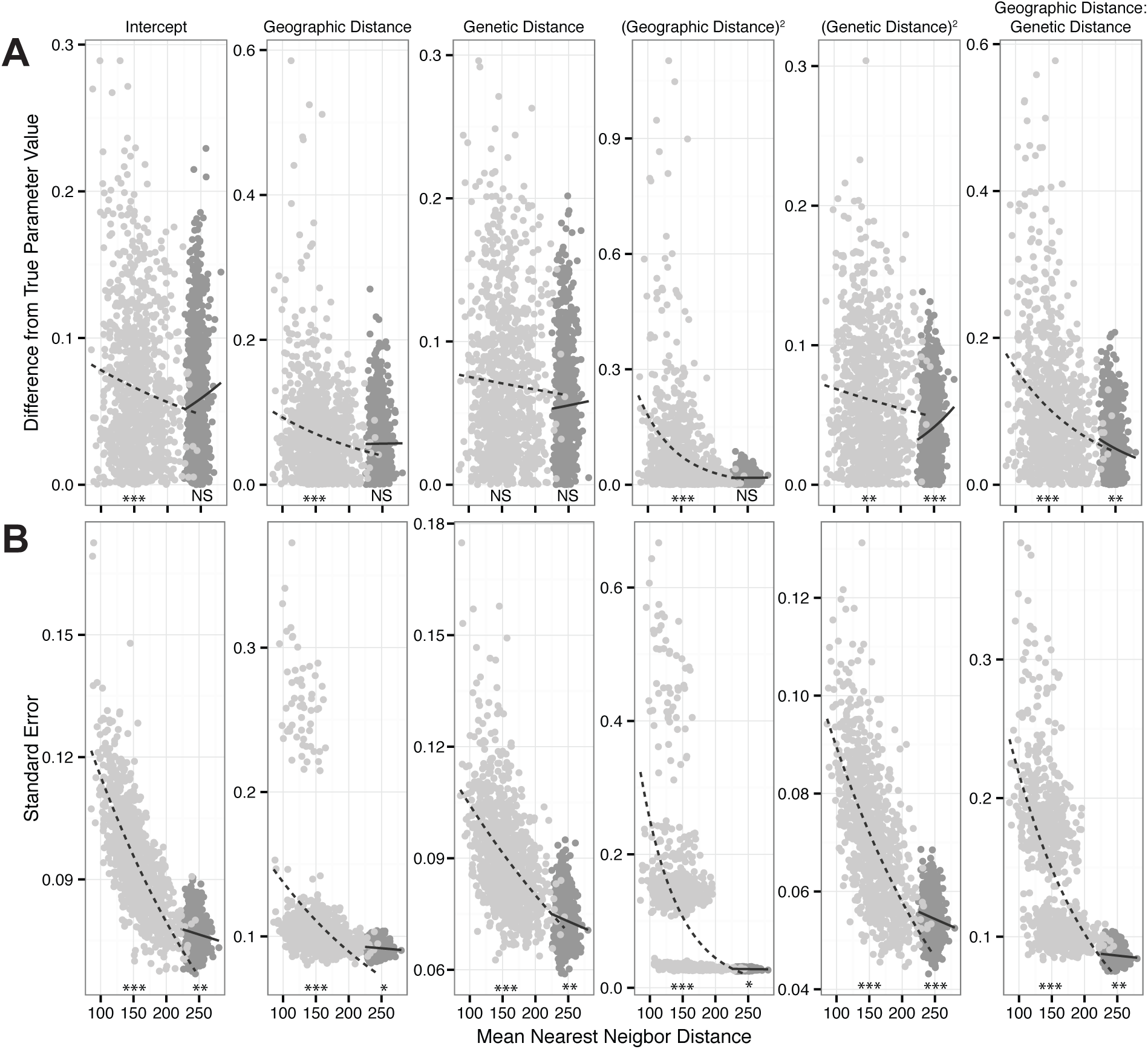
Relationship between measures of lack of model fit and the MNND of those crossing-sets for both SRS (light gray dots and dashed lines) and SPREAD-generated (dark gray dots and solid lines) crossing-sets. Trend lines were calculated using a generalized linear model with a Gaussian response distribution and a log link function. Significance of the line slope is shown below each line: *** p = 0, ** p=.001, * p =.01, NS p > .05. Panel **A**: The absolute value of the difference between “true” parameter values and parameter values of SRS- and SPREAD-generated crossing-sets for all six model parameters. Panel **B**: The standard errors of SRS- and SPREAD-generated crossing-sets for all six model parameters.

## DISCUSSION

SPREAD is an easily implemented algorithm designed to identify maximally informative, full factorial crossing-sets for use in mating experiments. SPREAD increases the diversity inherent in a crossing-set, for example, the genetic and geographic distances among crosses compared to a randomly sampled crossing-set. SPREAD requires two input parameters chosen by the user: the dimensions, *s*_*x*_ × *s*_*y*_, of the desired crossing-set and the number of randomly generated crossing-sets, *h*, from which the crossing-set with the MMNND is selected. SPREAD was designed for two dimensional trait data. If potential crosses are characterized by more than two target traits, and the traits are not completely independent, principal components analysis (PCA) can be used before implementing SPREAD to determine which two traits explain most of the trait variance (King and Jackson, 1999).

In our worked example, we successfully used SPREAD to select a crossing-set of 12 mat-A × 12 mat-a *N. crassa* strains. When these crosses are plotted in genetic vs. geographic distance space, it is evident that the selected set fulfills the desired criteria of evenly and completely covering the range of the larger set (Figure 1). Using the MMNND as the diversity metric favors crosses that are at the extremes of the trait-space. The inclusion of crosses with extreme trait distances in an original population should be carefully considered because these crosses will often be selected by SPREAD.

Calculating the true MMNND by computing MNND values for every possible subset of a sampled population may not be possible. Instead, our algorithm generates a large number (*h*) of random crossing-sets and chooses the set with the MMNND from those *h* crossing-sets. Choosing an appropriate *h* value is an important consideration when implementing the algorithm because too small an *h* value may reduce the probability of selecting a crossing-set with an MNND value close to the maximum value while too large an *h* value would unnecessarily increase computation time.

Our results show that the range of the distribution of MMNND values decreases as *h* value increases, mainly driven by an increase in the minimum MMNND value. Increasing the *h* value used in the algorithm increases the probability of obtaining an MMNND value close to the true MMNND value. However, the probability that an additional sample will yield a value higher than all samples already considered decreases as the number of samples already considered goes up. In terms of the number of random samples already sampled (*h*), this probability can be 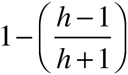 calculated as. For example, in our test, we used *h* values of 10, 100, 1000, and 10000. The probability of choosing an additional sample with a MNND value outside the range of the samples already taken would be .19, .0199, .001998, and 0.00019998, respectively. Increasing the *h* value above 1000 will result in a greater MMNND only 0.2% of the time. Based on this reasoning and our analyses of different *h* values, using an *h*-value of 1000 should be sufficient for most experiments with modest population sizes.

In experiments with a large breeding stock and small desired experimental mating population, there may be large variation in MNND values among crossing sets. Such large variation may offset the ability of even large *h* values to return a crossing-set that is substantially different from a randomly sampled crossing-set. In these cases, SPREAD could be modified to include a simulated annealing function that searches the space of a very large number of potential crossing-sets for a crossing-set that converges on a peak MNND value. One example of a simulated annealing algorithm that could be adapted for this purpose is SAGE (Simulated Annealing Guided Evaluation), developed to design combinatorial drug libraries (Zheng *et al.*, 1999).

We hypothesized that maximizing the diversity inherent in a crossing-set would increase the predictive ability of models relating outcomes to characteristics of crosses. When we compared model fits from crossing-sets generated by SPREAD to model fits from crossing-sets generated by SRS, we found that the model parameter values from SPREAD-generated crossing-sets were closer to the true model parameter values with smaller errors (Figure 3).

Although the parameter values from SPREAD-generated crossing-sets were closer to the true parameter values they did not precisely match the true parameter values. This is probably because the true parameter values are from a model calculated using the entire set of 24 mat-A × 24 mat-a crosses while the tested parameter values are from crossing-sets of 12 mat-A × 12 mat-a strains. The smaller sample size used to fit the model decreases both the precision and accuracy of estimated parameter values. Generalized linear models have been shown to be especially sensitive to sample size, compared to other methods (Wisz *et al.*, 2008).

The true MMNND does not need to be determined to produce crossing-sets that provide accurate estimates of underlying population level parameter values. For most parameters, SPREAD-generated crossing-sets with large MNND values fall at the bottom of the curve describing the relationship between the MNND of SRS-generated crossing-sets and either their deviation from true parameter values (Figure 4A), or the standard error of parameters (Figure 4B). Within the SPREAD-generated crossing-sets (dark gray, Figure 4), the benefit of a slightly higher MNND value to measures of model fit is negligible and, therefore, so is the benefit of increasing the *h* value used in SPREAD. Furthermore, the negative trend between the MNND of crossing-sets and our two measures of lack of model fit provides strong statistical support for our claim that maximizing the MNND of crossing-sets increases the utility of data generated from those crossing-sets.

Although our analysis used a dataset generated from the fungus *Neurospora crassa*, SPREAD can be easily used for many other organisms. SPREAD can be used to design breeding experiments for any fungus, plant, animal, or other sexual eukaryote with two mating-types or sexes where ecological, genetic, or physiological trait differences between the individuals involved in a cross are measured. The algorithm can be used directly, and does not require any modification when used with other organisms that satisfy these criteria.

For a species with more than two mating types, for example the social amoebae *Dictyostelium discoideum* that has three mating types (I, II, and III) (Bloomfield *et al.*, 2010), SPREAD would have to be run for each possible combinations of compatible mating types. The desired crossing-set size would have a maximum size dependent on the mating type with the fewest strains. For *D. discoideum*, SPREAD would have to be run six times for the following combinations: I×(II,III); II×(I,III); III×(I,II); I×II; I×III; II×III. The crossing set with the greatest MNND value from the six SPREAD selected crossing sets would then be used for experimentation. The number of SPREAD runs required will scale rapidly with the number of mating types involved, requiring parallel computation of SPREAD for species with large numbers of mating types.

SPREAD increases the value of fully crossed mating designs by enabling exploration and prediction across the full space of cross characteristics provided by available breeding stock. Simulations based on crossing-sets generated from the SPREAD algorithm *versus* SRS prove our algorithm generates more accurate parameter estimates, enabling better predictions of relationships between cross characteristics (*e.g.*, the genetic and geographic distances between parents) and the success of a cross. SPREAD is not computationally intense and is easy to implement, making it a valuable tool for researchers designing crossing experiments.

## ACKNOWLEDGEMENTS

We thank Kareem Carr and Steven Worthington for assistance in designing statistical analyses. We are grateful to members of the Pringle lab for advice and discussion. Our work is supported by the National Science Foundation Graduate Research Fellowship under Grant Nos. (DGE0644491 and DGE1144152) awarded to K.Z. and by other National Science Foundation grants awarded to the Pringle Laboratory. This work was also supported by funds from the Max Planck Institute for Demographic Research to K.Z., D.L., and A.P. Some computations in this paper were run on the Odyssey cluster supported by the FAS Division of Science, Research Computing Group at Harvard University. The work of D.L. was supported by the Max-Planck Odense Center, a collaboration of the Max Planck Society and the University of Southern Denmark.

